# Predicting Degradation Potential of Protein Targeting Chimeras

**DOI:** 10.1101/2024.09.16.613208

**Authors:** Andreas Petrou, Fayyaz Minhas

## Abstract

PRoteolysis TArgeting Chimeras (PROTACs) can inhibit protein activity by utilizing natural proteasomal degradation pathways for the degradation of target proteins. Being able to determine the degradation potential of PROTACs is crucial in drug development as it can lead to time, labor and cost savings. In this paper, we present a novel machine-learning pipeline that utilizes common compound fingerprints and a pre-trained graph neural network for the prediction of half-maximal degradation concentration of PROTACs by benchmarking a variety of protein tertiary structures and chemical features. Based on critical analysis of our cross-validation and independent test results, we have highlighted several key challenges underlying this prediction problem that need to be addressed to improve the generalization of predictive models in this domain. Moreover, we demonstrate the effectiveness of our approach by testing it on two different datasets and show that it performs better than the current state of the art with an AUC-ROC of 0.85 and accuracy of 0.875 on the DeepPROTACs test dataset.

## Introduction

In recent years, PRoteolysis TArgeting Chimeras (PROTACs) have emerged as a popular method for Targeted Protein Degradation (TPD). Common occupancy-driven molecules, used for regulating harmful protein function, are not always appropriate due to the lack of a binding site between the disease-causing protein and the drug. ^1^ The event-driven mechanism of action (MOA) of heterobifunctional drugs, such as PROTACs, allows for the removal of undruggable proteins from the system by utilizing proteasomal degradation as outlined in Figure 1.

**Figure 1.**
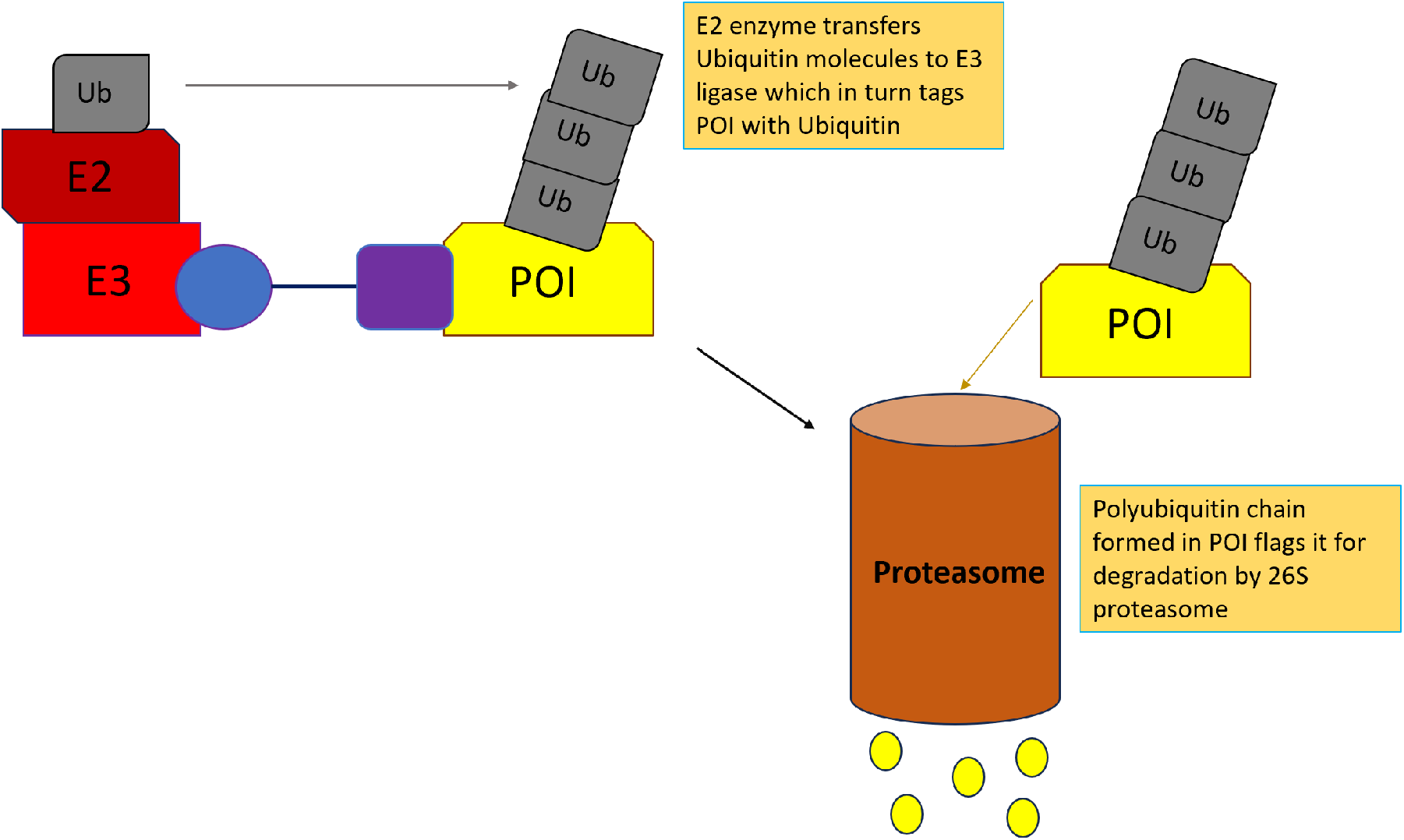
The E3 ligase, with the assistance of E2 enzymes, tags the target protein with Ubiquitin molecules, thus flagging the protein for destruction by the 26S proteasome. Then the proteasome takes the Ubiquitin-tagged protein and breaks it down into residues.

These types of drugs utilize the human body’s natural protein degradation system of mutated or misfolded proteins. E3 enzymes identify proteins that need to be removed from the system and tag them with Ubiquitin molecules, thus flagging them for destruction.^2,3^ A PROTAC is a ternary structure that consists of a ligand to a target protein of interest (POI), an E3 recruiting moiety that binds to the E3 ligase, and a linker that connects the protein and E3 ligands. Once a trimer of an E3 ligase, a PROTAC, and the POI is formed, E3 ligase tags the protein with ubiquitin molecules which trigger the proteasomal degradation pathway,^4^.^5^

As PROTACs advanced to Phase 1 and Phase 2 clinical trials, with ARV-110 administered to patients suffering from metastatic castration-resistant prostate cancer and ARV-471 administered to patients with metastatic breast cancer,^6^ the interest in these artificial protein degraders has surged. Nevertheless, the functionality of PROTACs depends on multiple steps and it is not sufficient for the drug to connect an E3 ubiquitin ligase to a target protein. ^7^ Binding affinity metrics such as inhibition constant (*K*_*i*_) and dissociation constant (*K*_*d*_)^8^ are not appropriate for estimating the effectiveness of PROTACs as binding does not guarantee degradation. It is not uncommon for companies to waste effort in producing thousands of candidate PROTAC compounds that are not capable of efficiently degrading proteins. Therefore being able to estimate the degrading capacity of these drugs can reduce labor and cost of production. Various measures exist for quantifying the effectiveness of PROTACs. Half-maximal degradation concentration (*DC*_50_) and maximal degradation (*D*_*max*_) are some of the measures used to classify whether a PROTAC is a good or bad degrader. A neural network, DeepPROTACs,^9^ used Graph Convolutional Neural Networks (GCNs) to extract features of molecules to classify PROTACs into ‘bad’ and ‘good’ degraders based on their *DC*_50_ and *D*_*max*_ values. Their network architecture is trained on a classification task and presents cross-validation results over a dataset derived from PROTACs-DB.^10^

The main focus of this paper is to find an appropriate regression model for finding the actual *DC*_50_ value of a candidate PROTAC. We have evaluated different machine learning models and feature representations for the prediction of *DC*_50_ of PROTAC candidates and identified significant practical limitations in the design of machine learning models in this domain.

## Methods

Fig 2 shows the overview of the proposed approach. Below, we discuss various steps involved in the development and evaluation of the proposed method.

**Figure 2.**
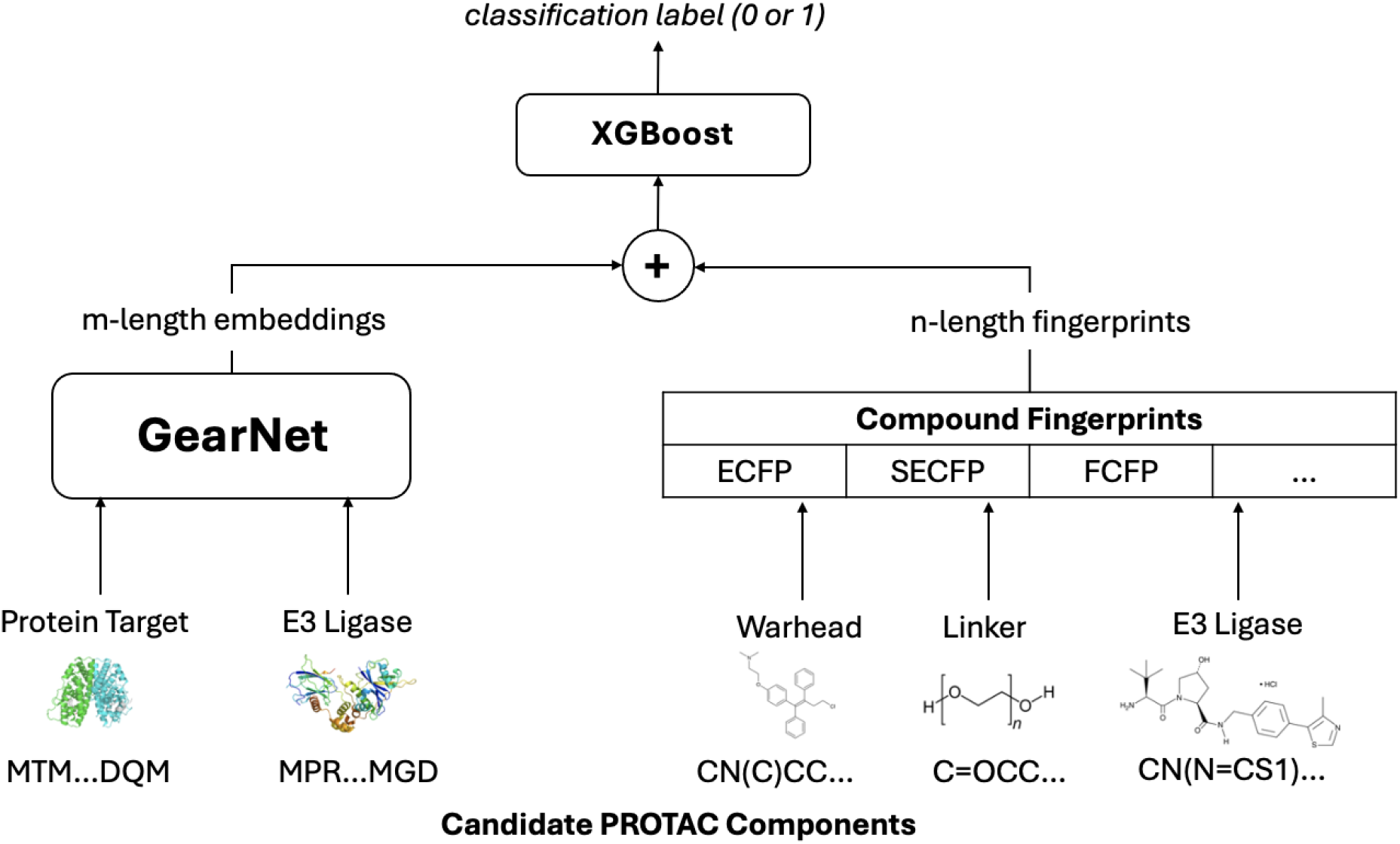
Model outline. PDB files of the proteins under investigation are used to generate their graph representations. Then these graph representations of the POI and E3 ligase are passed to a pre-trained GearNet for feature extraction while fingerprints are collected from the SMILES of PROTAC’s components. Then individual features are concatenated and the concatenated vector is passed to a Machine Learning model (XGBoost, random forest, or MLP) to predict the *DC*_50_ value of the PROTAC. Multi-format labels of individual examples allowed for the training of the models under investigation in both regression and classification tasks.

### Dataset

#### Databases

We have used PROTAC DB^10^ combined with PROTACpedia^11^ for model training and evaluation. Both databases contain information about PROTACs’ SMILES along with POI and E3 targets. They also include several drug properties such as *DC*_50_ and *D*_*max*_ along with the assays used to obtain these data. Overlapping examples found in both databases were removed to avoid inflated accuracy. In Table 1 we list the number of unique examples of the different entities (POI, E3 ligase, POI ligand, linker and E3 ligand) used in our prediction algorithm.

**Table 1:** The number of unique entities from PROTAC DB and PROTACPedia combined.

| ENTITY | NO. OF UNIQUE EXAMPLES |
| --- | --- |
| TARGET PROTEIN | 285 |
| E3 LIGASE | 17 |
| WARHEAD | 719 |
| LINKER | 1822 |
| E3 LIGAND | 182 |

The curated dataset we used for predicting the *DC*_50_ values of candidate PROTACs was obtained by collecting the SMILES of the PROTACs’ warheads, linkers, and E3 ligands along with the logarithm of their *DC*_50_ from the combined PROTAC database. The sequences of the protein of interest (POI) and the E3 protein were also obtained for each PROTAC.

### Feature Extraction

#### Graph-based Protein Features

Over the years, self-supervised methods and more specifically self-prediction models such as Bidirectional Encoder Representations from Transformers (BERT) provide a pipeline for pretraining text data for unsupervised tasks like language inference and answering questions. ^12^ Another language model is GPT-2 which is essentially a transformer implementation with layer normalization on self-attention block. ^13^ Both of these models performed equally well or sometimes even outperformed state-of-the-art models on a number of Natural language Processing Tasks. The success of these self-learning models encouraged researchers to pursue similar approaches to protein data as well. One such example is the Geometry Aware Relational Graph Neural Network.^14^ Graph networks can be more expressive than sequential data in the sense that they capture information such as protein active sites. This kind of data is impossible to obtain with the protein primary structures since sequences do not take into consideration protein shapes. Various versions of pre-trained GearNet exist based on residue type, distance, and bond angle. The protein embeddings used for training and testing were based on the distance loss function defined as 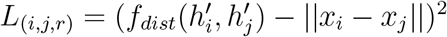.

Various versions of pre-trained GearNet exist where the graph neural network was trained on thousands of proteins from the AlphaFold database. Some of the loss functions used for training include multiview-contrastive learning, residue type prediction, distance prediction, dihedral prediction and angle prediction. Given the success of self-prediction methods in domains such as natural language processing, the version of Gearnet that is based on distance prediction was used to get embeddings from the proteins in our dataset.

GearNet takes a protein graph with residues as nodes, and constructs edges based on the sequential distance between nodes, K-nearest neighbor, and radius. The additional edges, required to construct the protein graph, increase memory requirements sufficiently, with a single V100 GPU requiring several hours to produce protein embeddings for the whole training set. The model weights were obtained through training with a cluster of four A100 GPUs with each GPU having a batch size of 2 (8 examples in each forward pass). We used a GearNet with 21 input dimensions (20 for amino acids and 1 additional dimension for padding), six layers, and a ReLU activation function along with batch normalization. Protein embeddings of the POI and E3 Ligase were generated, using model weights of a pre-trained GearNet on the AlphaFold protein database.^15^ These embeddings were then concatenated with compound fingerprints and passed to an XGBoost model for *DC*_50_ prediction.

#### Compound Representations

For the warhead, linker and E3 ligand, we used various methods for extracting features from their SMILES. Molecular fingerprints such as Morgan, which is also known as Extended-Connectivity Fingerprint or ECFP, are fixed-length vectors appended with ones and zeroes based on the substructures present within a molecule.

Over the years there have been numerous methods for featurizing SMILES. However, there is no optimal SMILES representation yet. This encouraged us to experiment with different types of molecular representations ^16^ to analyze how different fingerprints affect the predictive capabilities of the models under consideration. These include structural fingerprints (such as ECFP, SMILES Extended Connectivity Fingerprint (SECFP) and Molecular Access System (MACCS)) and physico-chemical descriptors such as 2D RDKit descriptors.

### Predictive Models and evaluation

Based on the various feature representations discussed earlier, we used an XGBoost predictor to output the final value, denoting the degradation capacity of the candidate drug under investigation. XGBoost stands for extreme gradient boosting and it is an ensemble model based on decision trees. It uses successive trees to try and ‘correct’ model prediction and have produced excellent results on various complex machine-learning tasks such as image recognition. In addition, they are convenient when the dimension of the feature vectors is greater than the number of examples,^17^ hence were rendered suitable for generating predictions out of our extracted features.

In the following sections, we discuss how our proposed algorithm fare against our prediction task and compare its performance against the current state of the art, DeepPROTACs, based on classification metrics including Area Under the ROC curve (AUC-ROC) and accuracy.

### External validation

For tuning our models and selecting an appropriate fingerprint extraction method we used the CD-HIT^18^ algorithm on target proteins to produce folds based on protein similarity. CD-HIT is an optimized sequence alignment procedure that utilizes multi-processing to compute the similarity between long chains of amino acids in a reasonable amount of time. The folds, generated from CD-HIT clustering, were used to split the data into 10 parts which were further used to separate the dataset into train and validation with a 9:1 ratio.

Even though our dataset used for training has the actual *DC*_50_ value of the candidate PROTACs, this information is not available on the two datasets used for testing. To solve this issue we trained our proposed regression models on continuous labels and tested them on binary ones. Continuous labels were converted to binary values (1 or 0) based on the criteria given on.^9^

Similar to the existing DeepPROTACs classification, ^9^ we also tested the predictors from our validation analysis on an independent test consisting of 16 PROTACs aimed at targeting estrogen receptor (see Table 2). Limiting estrogen receptor (ER) activity has become one of the main focuses for treating breast cancer. ^19^ ER ligands, such as toremifene, are common drugs for treating this kind of cancer. Therefore, having a drug capable of degrading ER through targeted proteasomal degradation can prove very useful. Given this, we tested our models on ER degrading PROTACs and although in each example, the linker is altered slightly, this has a significant impact on the degrading capabilities of the artificial ER degraders. The PROTACs in the test dataset have the same warhead (toremifene) and E3 ligand (VHL032) while the linkers, which are based on Alkyl and PEG chains, have an additional Carbon atom between successive examples. As only categorical *DC*_50_ values (high or low) are available as ground truth for this set, we used the regression model as a classifier to compare its prediction accuracy and AUC-ROC to results reported in.^9^

**Table 2:**
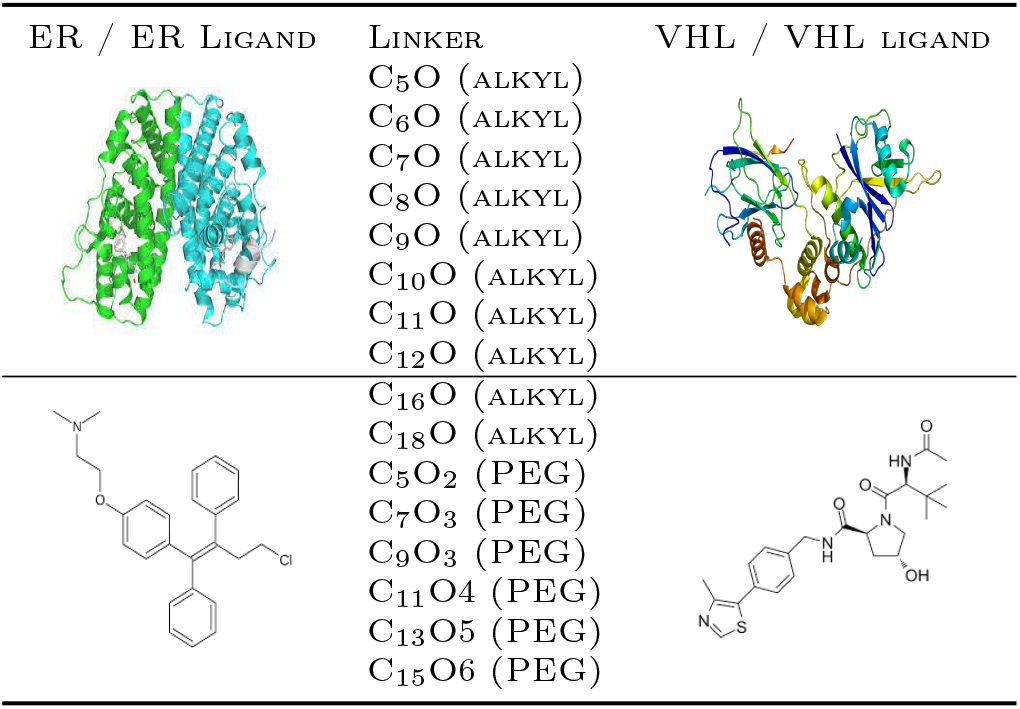
ER degrading PROTACs in the DeepPROTACs independent test set.

| ER / ER LIGAND | LINKER | VHL / VHL LIGAND |
| --- | --- | --- |
|  | C<sub>5</sub>O (ALKYL) |  |
|  | C<sub>6</sub>O (ALKYL) |  |
|  | C<sub>7</sub>O (ALKYL) |  |
|  | C<sub>8</sub>O (ALKYL) |  |
|  | C<sub>9</sub>O (ALKYL) |  |
|  | C<sub>10</sub>O (ALKYL) |  |
|  | C<sub>11</sub>O (ALKYL) |  |
|  | C<sub>12</sub>O (ALKYL) |  |
|  | C<sub>16</sub>O (ALKYL) |  |
|  | C<sub>18</sub>O (ALKYL) |  |
|  | C<sub>5</sub>O<sub>2</sub> (PEG) |  |
|  | C<sub>7</sub>O<sub>3</sub> (PEG) |  |
|  | C<sub>9</sub>O<sub>3</sub> (PEG) |  |
|  | C<sub>11</sub>O<sub>4</sub> (PEG) |  |
|  | C<sub>13</sub>O<sub>5</sub> (PEG) |  |
|  | C<sub>15</sub>O<sub>6</sub> (PEG) |  |

Apart from the independent test dataset used to test DeepPROTACs, we also constructed our dataset, with a variety of target proteins aiming at testing how well our proposed models cope with unseen examples. The new test examples include candidate PROTACs that attempt to degrade proteins involved in diseases with high mortality rates such as breast cancer and prostatic cancer.

Our new test dataset has a similar structure to the DeepPROTACs independent test set and our training dataset, constructed from PROTAC DB and PROTACPedia. It includes two proteins, a POI and an E3 ligase, along with the components of the candidate PROTAC which are the linker, warhead and E3 ligand. Some of the candidate drugs included in the custom dataset include ER and its derivatives such as estrogen-related receptor alpha denoted as *ERRα* whose activity lead to breast cancer. Other proteins, with harmful activities, that are targetted by candidate PROTACs in our test dataset include Focal Adhesion Kinase (FAK) and TANK-binding kinase (TBK 1) linked to tumor microenvironments and pancreatic cancer respectively.

## Results and Discussion

In this section, we present the performance of our proposed models that utilize GearNet embeddings and common molecule fingerprints to predict the degradation potential of candidate PROTACs. We also showcase the effectiveness of our optimal model by comparing it against the current state of the art, which is DeepPROTACs.

### Validation results

As discussed earlier, our model pipeline was trained on the regression task of predicting the actual *DC*_50_ value of candidate PROTACs and then tested on classifying potential protein degraders into ‘good’ and ‘bad’. Although our models were trained on regression labels, all results reported used classification metrics as these metrics were used to estimate the performance of DeepPROTACs as well.

For fine-tuning the machine learning models, we split the dataset into training and validation by using the protein clusters produced from CD-HIT based on the similarity of target proteins. Then we used these clusters to create train and validation sets with a 9:1 ratio.

From the results reported in Table 3, it can be observed that XGBoost provides good cross-validation results with high AUC-ROC achieved by all different fingerprint combinations indicating that our model is not unbiased towards the positive or negative class. In addition, the fingerprint that produced the best results was AVALON achieving an AUC-ROC of 0.90 and accuracy of 0.73. Although LAYERED and SECFP achieved same AUC-ROC as AVALON they achieved less accuracy at 0.67 and 0.66 respectively.

**Table 3:**
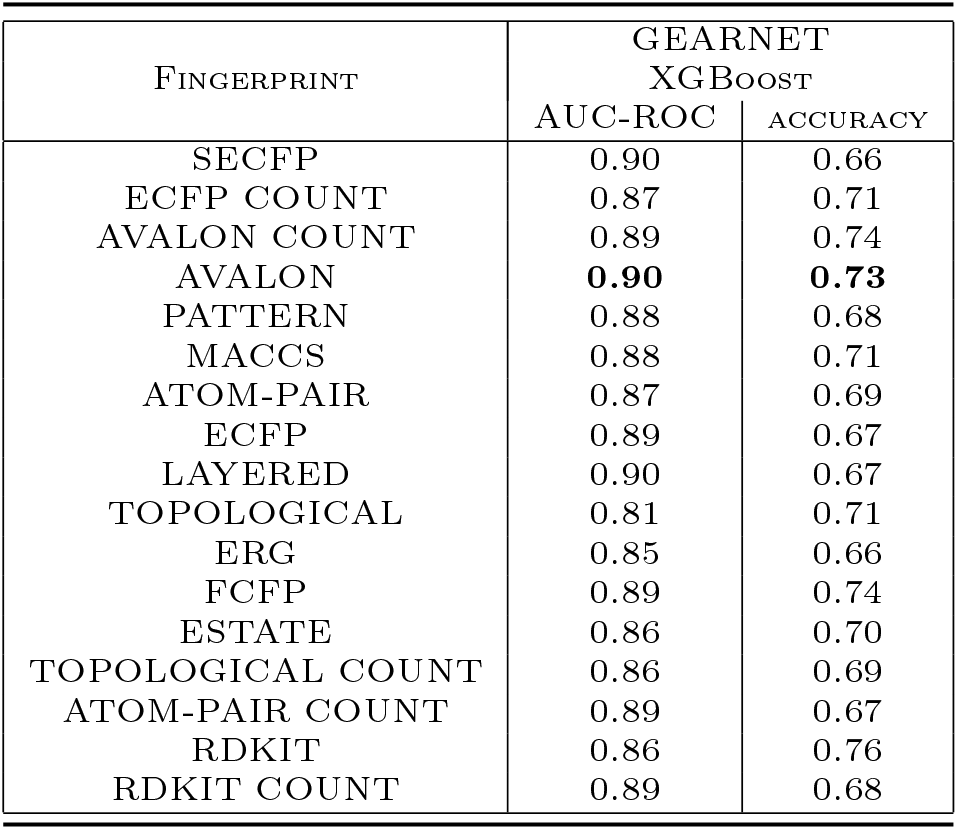
Validation results of XGBoost on the classification task of splitting candidate PROTACs into ‘good’ and ‘bad’ degraders. The dataset was split on train and validation with a 9:1 ratio using the CD-HIT algorithm to divide the dataset into clusters based on the similarity of target proteins. Models were evaluated using AUC-ROC and accuracy.

### Testing on unseen data and comparison

For testing, we used two different datasets. The dataset on the ER-degraders with different linkers used by Li et. al^9^ and our custom dataset composed of recently discovered PROTACs targetting a variety of target proteins not included during training.

The DeepPROTACs test dataset has 16 distinct examples which are not evenly split among the two classes (positive and negative) with 11 positive examples and 5 negative ones. As a result, for a model to achieve satisfactory results on this test dataset it is necessary to attain both high accuracy and AUC-ROC.

The mismatch between positive and negative examples led to a discrepancy between accuracy and AUC-ROC as it can be observed in Table 4. Based on the validation results the optimal fingerprint for XGBoost (AVALON) outperformed DeepPROTACs with an accuracy of 0.875 compared to DeepPROTACs’ accuracy of 0.6875. It also managed to score an AUC-ROC of 0.85 demonstrating that it is unbiased towards any of the two classes in the DeepPROTACs test dataset. In addition, AVALON fingerprints achieved similar results with the custom test dataset as well providing an AUC-ROC of 0.88 and an accuracy of 0.85 indicating good generalization on both test datasets.

**Table 4:** Test results of XGBoost model trained on a dataset with 9:1 ratio train-validation split obtained by using CD-HIT to produce clusters of examples based on similarity of the target protein. Best performant fingerprints on the validation set have their test performance highlighted in bold. Model evaluation on the DeepPROTACs test dataset and our custom dataset is provided through the AUC-ROC and accuracy achieved by the different models.

| FINGERPRINT | DEEPPROTACs TEST |  | CUSTOM DATASET |  |
| --- | --- | --- | --- | --- |
|  | XGBoost |  |  |  |
|  | AUC-ROC | ACCURACY | AUC-ROC | ACCURACY |
| SECFP | 0.5 | 0.6875 | 0.88 | 0.89 |
| ECFP COUNT | 0.5 | 0.6875 | 0.82 | 0.77 |
| AVALON COUNT | 0.8 | 0.75 | 0.82 | 0.89 |
| AVALON | **0.85** | **0.875** | **0.88** | **0.85** |
| PATTERN | 0.38 | 0.6875 | 0.75 | 0.77 |
| MACCS | 0.75 | 0.8125 | 0.76 | 0.69 |
| ATOM-PAIR | 0.45 | 0.625 | 0.78 | 0.72 |
| ECFP | 0.5 | 0.6875 | 0.81 | 0.87 |
| LAYERED | 0.33 | 0.6875 | 0.75 | 0.79 |
| TOPOLOGICAL | 0.35 | 0.6875 | 0.89 | 0.9 |
| ERG | 0.53 | 0.5 | 0.73 | 0.62 |
| FCFP | 0.5 | 0.6875 | 0.85 | 0.85 |
| ESTATE | 0.62 | 0.75 | 0.48 | 0.79 |
| TOPOLOGICAL COUNT | 0.5 | 0.625 | 0.90 | 0.82 |
| ATOM-PAIR COUNT | 0.6 | 0.75 | 0.67 | 0.69 |
| RDKit | 0.61 | 0.6875 | 0.70 | 0.79 |
| RDKit COUNT | 0.48 | 0.625 | 0.79 | 0.82 |

### Critical analysis of results

Given the limited number of labeled data, it comes as no surprise that our traditional machine-learning methods surpass the classification performance of neural networks^20^ or more complicated pipelines.

Although the reported methods give good cross-validation performance and some of our test results surpass the existing state of the art, despite being simpler, a lot of our models falter in the test dataset with an AUC-ROC exactly or close to 0.50 suggesting that many of our models are biased towards one class over another in the DeepPROTACs test dataset. This can be due to the feature representations of chemical compounds, being inadequate in effectively capturing small changes in the drug structure. Even though it is evident from the data that changing the PROTAC linker can affect the potential drug’s degradation capacity, we noticed that the output of the prediction model undergoes a small change in its value when the linker is changed. This is probably due to the fact that adding more atoms to alkyl and PEG chains in the linker hardly changes their fingerprints. It can be concluded that using fingerprints for featurizing SMILES may not be the optimal option as slightly changing the linker can alter the degrading capacity of the candidate drug but not the representation of the linker molecule.

Another possible reason for the poor generalization of some of our models is that available PROTACs are not only small in number but also quite homogeneous in their chemical composition. Although PROTAC DB has a plethora of PROTACS and numerous linkers, warheads, and E3 ligands, the number of E3 ligases is fairly limited with only 8 significantly distinct E3 ligases in comparison to the 600 or so E3 ligases available in the human body. This can lead to the examples available in training and cross-validation being very similar to each other resulting in an overestimate of prediction performance when using classical cross-validation.

In order to test this hypothesis, we performed dimensionality reduction with principal component analysis (PCA) and clustering of the entire PROTACs DB and PROTACPedia datasets used in this work for training and validation. We observed three major clusters based on the similarity of E3 ligases (Figure 3). This leads us to believe that the high predictive performance in the original validation analysis in our work as well as the classification-based DeepPROTACs model is possibly due to the training and test examples being more similar than can be expected in a realistic test environment. Thus, these models may not be able to reliably predict the degradation potential of test PROTACs that have different E3 ligases and E3 targetting moieties in comparison to those used in training. A similar issue in the evaluation of machine learning models for protein-protein interactions has been reported in This is a fundamental challenge in this domain that requires further investigation and the design of improved pipelines before such methods can be used in practice.

**Figure 3.**
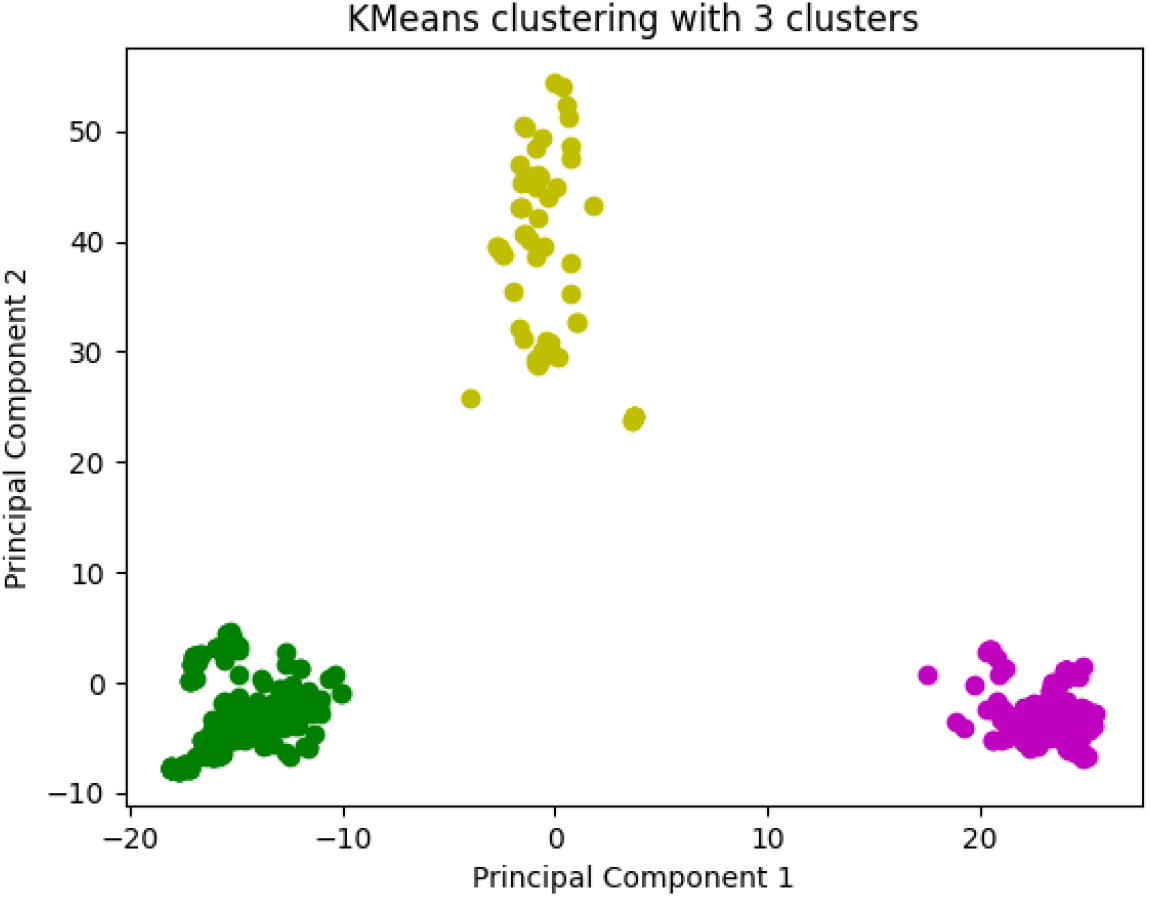
Principal Component Analysis on the training dataset shows 3 major clusters indicating significant homogeneity in terms of E3 ligases.

### Model deployment and code availability

Despite the challenges mentioned previously, some of our models managed to achieve commendable performance with some of them even surpassing the current state of the art; DeepPROTACs. Given this, we deployed our best model which is an XGBoost regressor with GearNet embeddings and AVALON fingerprints for researchers to use on their PROTACs for better insights into their effectiveness. Our model can be used at https://huggingface.co/spaces/AndreasP1999/PROTAC-DC50 and our code is available at https://github.com/MonkeyDLuffy99/Masters-FINAL

### Conclusions and Future Work

In this work, we have developed baseline methods for the prediction of the degradation potential of PROTACs. Our initial analysis shows that although the proposed regression model and existing classifiers can offer high cross-validation performance, further work is needed to improve their generalization performance. So far we have tested our proposed pipelines only on a small subset of potential drugs that can be used for targetted protein degradation. In the future, we plan to extend this work by the use of more effective feature representations, a larger training and test dataset as well as simultaneous prediction of multiple degradation measures such as IC50, Dmax, and percentage degradation.

